# SERPINE1–AP2A1 interplay links substrate stiffness to fibroblast senescence

**DOI:** 10.64898/2026.06.30.735455

**Authors:** Intan Chairun Nisa, Pirawan Chantachotikul, Takumi Saito, Cristina Bertocchi, Shinji Deguchi

## Abstract

Cellular senescence is characterized by stable cell-cycle arrest, cytoskeletal remodeling, and altered secretion of senescence-associated secretory phenotype (SASP) factors, including SERPINE1/plasminogen activator inhibitor-1 (PAI-1). Although extracellular matrix (ECM) stiffening has been linked to fibroblast mechanotransduction and SERPINE1-associated remodeling, the molecular pathway connecting substrate stiffness to SERPINE1 regulation in senescent fibroblasts remains incompletely understood. Here, we investigated how defined substrate stiffness affects fibroblast morphology, mechanical phenotype, and SERPINE1 expression, and examined whether the clathrin adaptor AP2A1 participates in this response in replicative senescent human fibroblasts. Using tunable polyacrylamide hydrogels, we found that increasing substrate stiffness enhanced fibroblast spreading, stress fiber thickening, focal adhesion maturation, cellular stiffness, and senescence-associated marker expression. Stiff substrates also increased SERPINE1 expression and its colocalization with actin fibers, with stronger responses observed in senescent than in young fibroblasts. Functional perturbation experiments further suggested that SERPINE1 contributes to stress fiber organization in senescent cells. In addition, AP2A1 colocalized with SERPINE1, and modulation of AP2A1 under knockdown and overexpression conditions altered SERPINE1 signal intensity. Conversely, perturbation of SERPINE1 also affected AP2A1, supporting a potential bidirectional relationship between these two components. Together, these findings identify SERPINE1 as a stiffness-responsive factor associated with senescence-linked cytoskeletal remodeling and support a functional relationship between AP2A1 and SERPINE1 in senescent fibroblasts. These results suggest that the AP2A1-SERPINE1 axis may contribute to the link between extracellular mechanical cues and senescence-associated fibroblast remodeling.

## Introduction

Cellular senescence is a state of irreversible cell cycle arrest triggered by diverse stressors, including telomere shortening [1], [2], [3], mitochondrial dysfunction [4], [5], activation of the DNA damage response [6], and epigenetic modifications [7], [8]. Senescent cells also undergo marked phenotypic remodeling, characterized by morphological enlargement, enhanced cell–substrate adhesion, cytoskeletal reorganization, and secretion of senescence-associated secretory phenotype (SASP) factors [9], [10], [11], [12], [13]. In fibroblasts, the development of hypertrophic actin stress fibers is a prominent feature of the senescent state and is associated with enhanced cell–substrate mechanical interactions [14], [15].

Recent proteomic analyses have revealed extensive remodeling of stress fiber composition in human fibroblasts during replicative senescence, accompanied by increased abundance of numerous stress fiber-associated proteins [14]. Several of these proteins have been implicated in the regulation of senescence-associated phenotypes and their partial reversibility at the cellular level [14], [15]. Among them, AP2A1 (alpha 1 adaptin subunit of the adaptor protein 2) has been shown to regulate integrin β1 trafficking to cell–substrate adhesions, thereby contributing to the structural stability of enlarged senescent cells [15]. Another stress fiber-associated protein enriched during fibroblast senescence is SERPINE1, which encodes plasminogen activator inhibitor-1 (PAI-1). SERPINE1 is widely recognized as both a biomarker and a functional mediator of senescence and has been implicated in extracellular matrix (ECM) remodeling, fibrosis, tissue stiffening, and maintenance of senescent phenotypes [16], [17], [18], [19]. While SERPINE1 is well established as a secreted mediator of senescence, its functional association with stress fiber remodeling and intracellular trafficking before extracellular release remains incompletely understood.

Mechanical cues from the extracellular microenvironment are important regulators of senescence. Age-associated ECM stiffening alters fibroblast morphology, promotes focal adhesion maturation, and reorganizes actin stress fibers, thereby reinforcing cellular tension and mechanotransduction [11], [14], [20]. Stiff substrates are also associated with increased SERPINE1 expression, suggesting that SERPINE1 may function as a mechanosensitive mediator of senescence-associated remodeling [21], [22]. Several pathways activated by matrix stiffness, including integrin-dependent signaling, RhoA/ROCK, and YAP/TAZ, have been implicated in linking extracellular mechanics to SERPINE1 regulation [20], [23]. However, how extracellular mechanical cues are integrated with cytoskeletal remodeling to regulate SERPINE1 trafficking, secretion, and function in senescent fibroblasts remains unclear.

Here, we examined how extracellular matrix stiffness shapes SERPINE1-associated remodeling in senescent fibroblasts, focusing on the stress fiber-associated adaptor protein AP2A1. We show that substrate stiffness enhances senescence-associated mechanophenotypes and promotes SERPINE1 expression and its association with stress fibers. We further identify a functional interplay between intracellular AP2A1 and the extracellular matrix regulator SERPINE1, accompanied by their coordinated movement along stress fibers. These findings define a molecular link between extracellular mechanical cues and SERPINE1-dependent remodeling, supporting the view that matrix stiffness actively modulates the senescent state rather than simply reflecting it.

**Figure 1.**
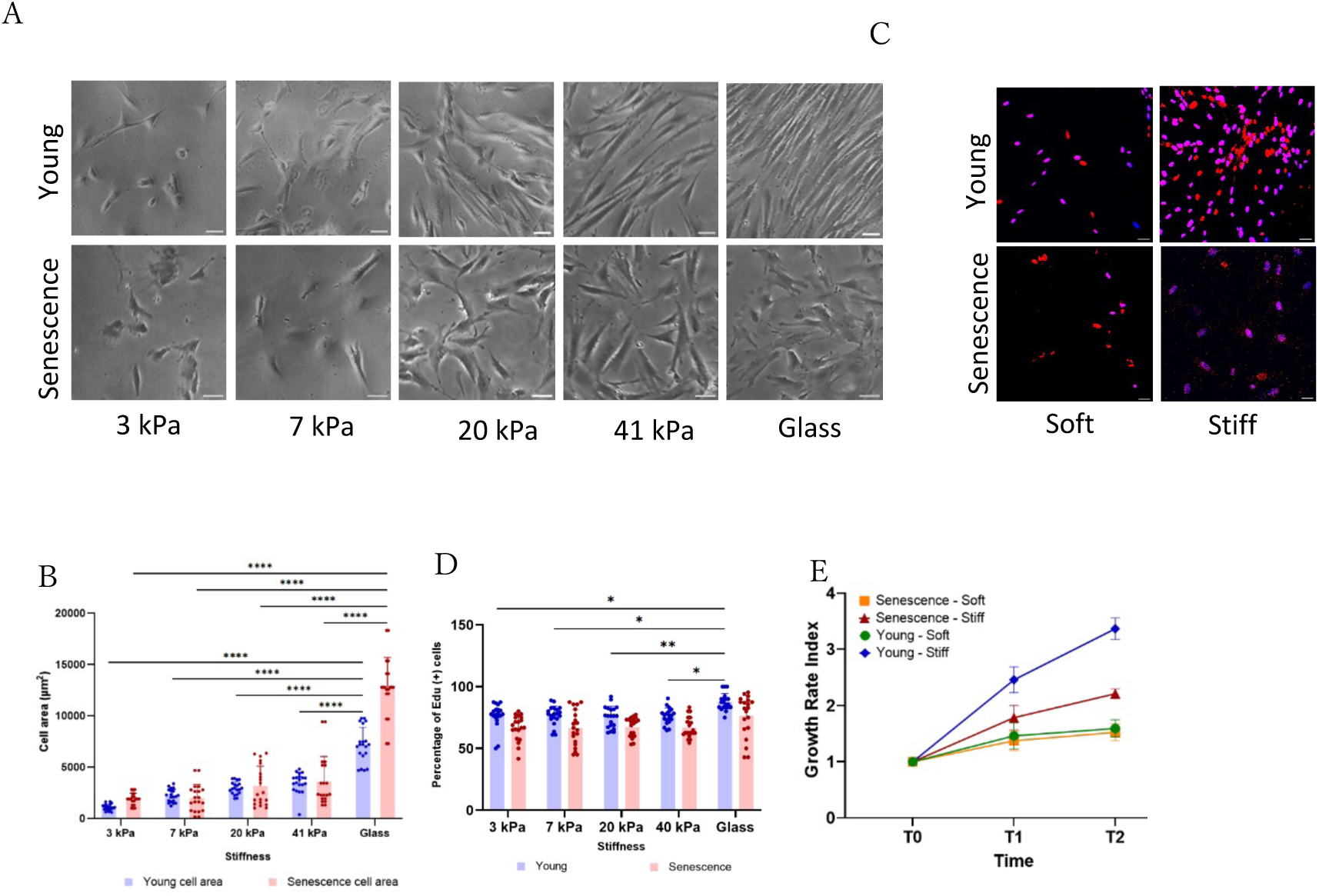
Substrate stiffness controls fibroblast morphology and proliferation. A) Representative images of HFF-1 fibroblasts cultured on polyacrylamide hydrogels of defined stiffness (3, 7, 20, 41 kPa) and glass, imaged by light microscopy at 4× magnification (Scale bar: 50 μm). B) Quantification of cell spreading area (n = 30 cells from three independent experiments). C) EdU-positive cells (magenta) indicating proliferation for each stiffness condition. Comparisons were made between soft (3 kPa) and stiff (glass) substrates (Scale bar: 50 μm). D) Percentage of cells positive for EdU staining, indicating proliferative activity (n = 30). E) Cell growth index measured at 24 h (T0), 48 h (T1), and 72 h (T2); values at T0 were used as baseline. Comparisons were made between soft substrates (3 kPa) and stiff substrates (glass).

## Results

### Substrate stiffness enhances senescence-associated mechanophenotypes

Human foreskin fibroblasts (HFF-1) were serially passaged to induce replicative senescence, with early-passage cells (P10) used as the young group and late-passage cells (P30) used as the senescent group. Cells were cultured on defined polyacrylamide hydrogels spanning physiological stiffnesses (3–41 kPa) relevant to soft tissues such as skin, as validated by atomic force microscopy (AFM) (Supplementary Fig. S1A,B).

Increasing substrate stiffness promoted stress fiber formation and thickening in both young and senescent cells, with a more pronounced response in senescent cells (Figures 2A–C). Quantitative analysis showed that the number of stress fibers increased with substrate stiffness, while senescent cells exhibited a higher fraction of thick stress fibers than young cells. A similar trend was observed for focal adhesions, in which paxillin staining revealed an increase in area with substrate stiffness in both cell populations. Senescent cells exhibited larger focal adhesions than young cells, particularly on stiffer substrates, although this increase appeared to plateau above 20 kPa (Supplementary Fig. S3F–H). AFM measurements further showed that cellular Young’s modulus increased with substrate stiffness in both young and senescent cells. Across the stiffness range, senescent fibroblasts exhibited higher cellular stiffness than young cells, in parallel with their more prominent stress fibers and focal adhesions (Supplementary Fig. S2F–I).

**Figure 2.**
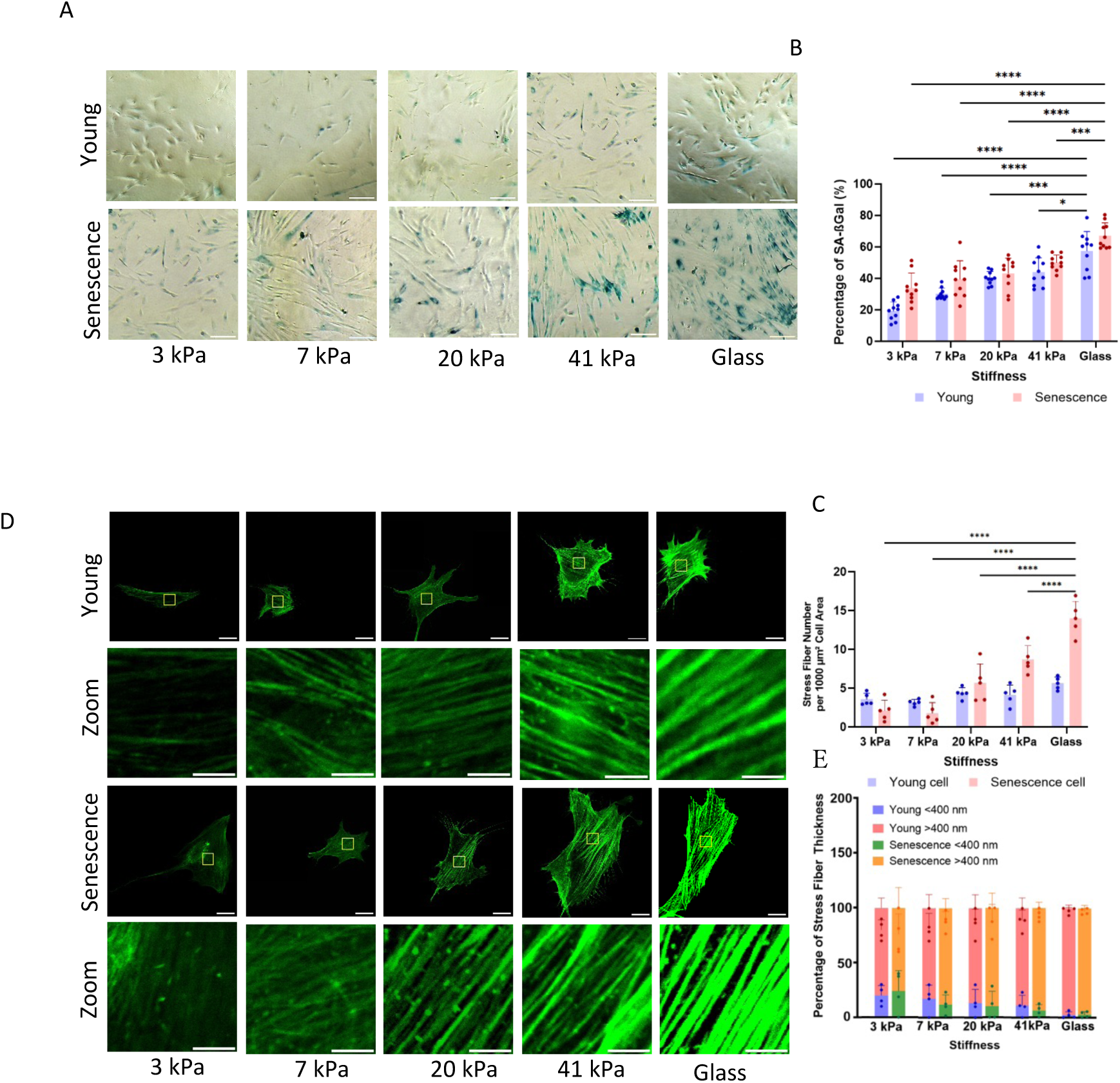
Substrate stiffness modulates senescence-associated cytoskeletal remodeling. A) Immunofluorescence staining for F-actin (phalloidin) in young and senescent fibroblasts cultured on substrates of varying stiffness (scale bar: 25 μm; inset: 2 μm). B) Quantification of stress fiber number per 1,000 μm² cell area (n = 30 cells). C) Percentage of thick stress fibers among all measured fibers (n = 300 fibers across 30 cells). D) Representative images of senescence-associated β-galactosidase (SA-β-gal) staining in fibroblasts cultured on substrates of varying stiffness. SA-β-gal-positive cells are shown in blue. E) Percentage of SA-β-gal-positive cells across substrate stiffness conditions (n = 20).

Consistent with these structural and mechanical changes, senescence-associated β-galactosidase (SA-β-gal) positivity increased with substrate stiffness (Figure 2D,E). Senescent cells showed higher baseline SA-β-gal positivity than young cells, which was further elevated on rigid substrates. In addition, p53 expression increased with substrate stiffness, as shown by both immunofluorescence and western blot analyses (Supplementary Fig. S3A–E). Together, these results show that substrate stiffness enhances multiple structural, mechanical, and molecular features associated with cellular senescence.

### Substrate stiffness induces SERPINE1 expression and stress fiber association

We next examined whether SERPINE1 expression was similarly regulated by substrate stiffness. Immunofluorescence analysis showed that SERPINE1 signal increased with substrate stiffness in both young and senescent cells, with senescent cells consistently exhibiting higher SERPINE1 signal than young cells across all conditions (Figure 3A, B).

**Figure 3.**
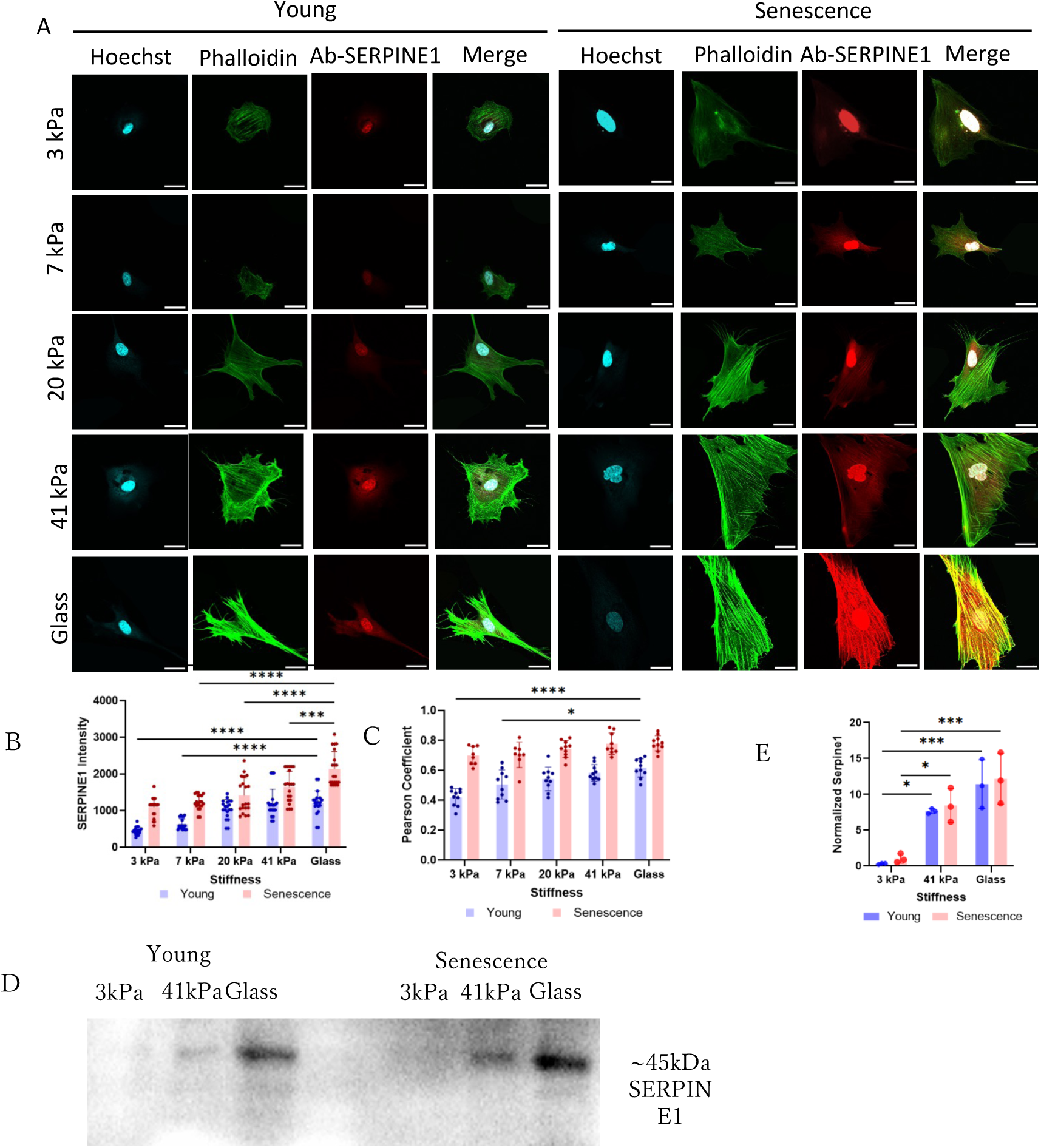
Substrate stiffness induces SERPINE1 expression in young and senescent fibroblasts. A) Representative immunofluorescence images of SERPINE1 staining in young and senescent fibroblasts cultured on substrates of varying stiffness (3, 7, 20, and 41 kPa, and glass). SERPINE1, red; F-actin/phalloidin, green; nuclei, blue (scale bar: 25 μm). B) Quantification of mean SERPINE1 fluorescence intensity (n = 30 cells per condition). C) Quantification of SERPINE1 colocalization with actin stress fibers (Pearson’s correlation coefficient). D) Western blot analysis of SERPINE1 in cell lysates from fibroblasts cultured on 3 kPa and 41 kPa hydrogels and glass coverslips. The band at ∼45 kDa corresponds to SERPINE1. E) Quantification of normalized SERPINE1 protein levels in young (P11) and senescent (P31) fibroblasts cultured on soft (3 kPa) and stiff (41 kPa) substrates.

To determine whether substrate stiffness altered the association between SERPINE1 and stress fibers, we used structured illumination microscopy (SIM) to quantify SERPINE1–actin colocalization across stiffness conditions. Pearson’s correlation coefficient increased progressively with substrate stiffness in both cell populations and remained higher in senescent fibroblasts than in young cells (Figure 3C). These results suggest that SERPINE1 becomes more closely associated with stress fiber-rich regions under stiffer conditions.

Western blot analysis further showed that SERPINE1 protein levels increased with substrate stiffness in fibroblasts cultured on 3 kPa, 41 kPa, and glass substrates (Figure 3D). Quantification confirmed higher SERPINE1 abundance in senescent cells than in young cells under both soft and stiff conditions (Figure 3E), consistent with the immunofluorescence analysis. This senescence-associated increase was supported by comparison of young and senescent cell lysates (Supplementary Fig. S4A–B). Together, these results show that SERPINE1 expression is increased by substrate stiffness and further enhanced in senescent fibroblasts. In addition, increased SERPINE1 expression was accompanied by enhanced colocalization with stress fibers.

### SERPINE1 regulates stress fiber organization in senescent fibroblasts

To determine whether SERPINE1 contributes to stress fiber organization, we manipulated SERPINE1 expression in passage-31 (P31) senescent fibroblasts. siRNA-mediated knockdown of SERPINE1 reduced intracellular SERPINE1 signal and resulted in a marked decrease in phalloidin-positive stress fibers compared with control cells (Figure 4A,B). In contrast, overexpression of SERPINE1 using an mClover-SERPINE1 construct increased SERPINE1 signal and stress fiber intensity (Figure 4C,D). Together, these results suggest that SERPINE1 positively regulates stress fiber organization in senescent fibroblasts.

**Figure 4.**
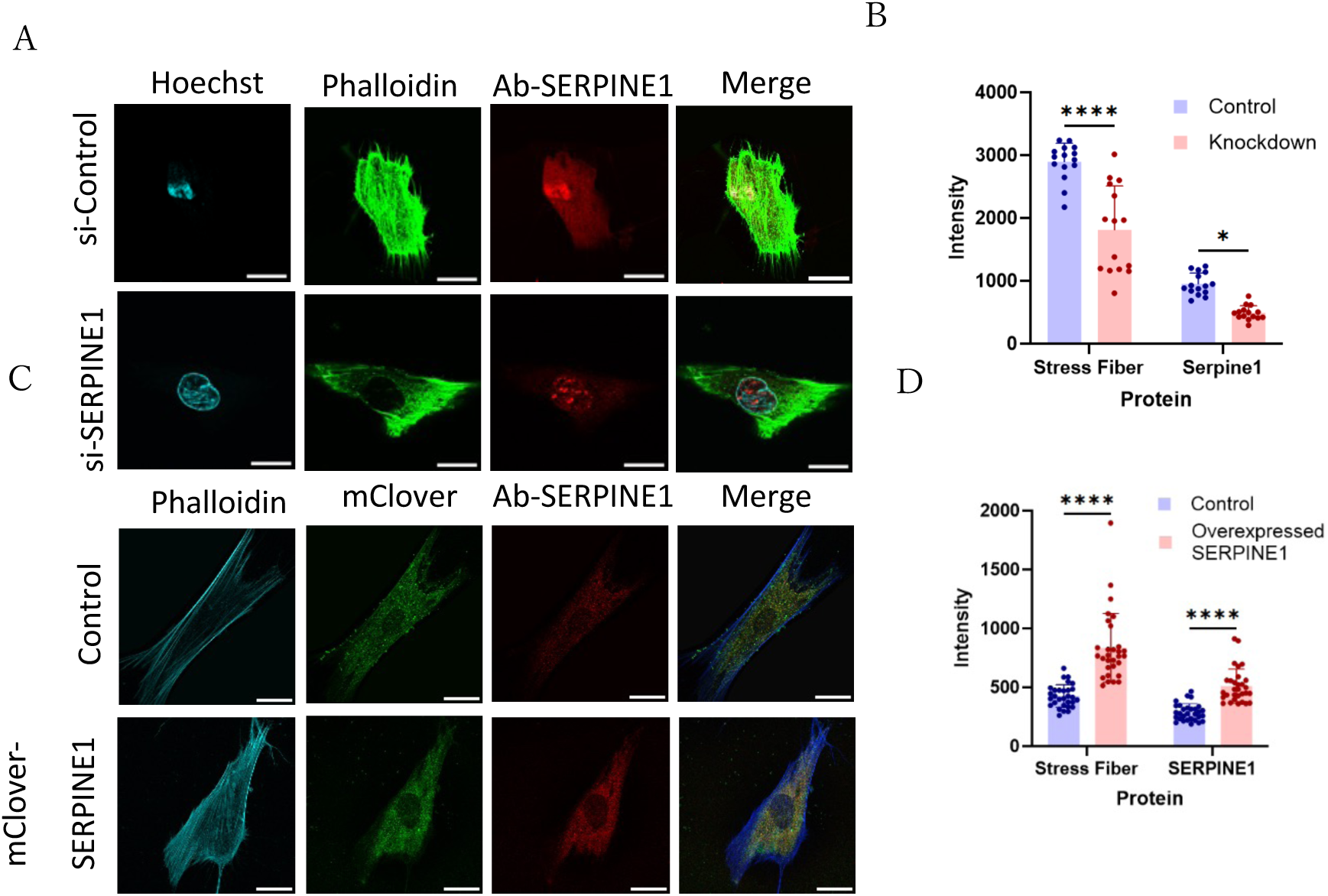
SERPINE1 regulates stress fiber remodeling in senescent fibroblasts. A) Representative immunofluorescence images of SERPINE1-knockdown cells stained for SERPINE1, F-actin/phalloidin, and nuclei (scale bar: 25 μm). B) Quantification of stress fiber intensity after SERPINE1 knockdown (n = 30 cells). C) Representative images of mClover-SERPINE1-overexpressing cells stained for SERPINE1 and F-actin/phalloidin (scale bar: 25 μm). D) Quantification of stress fiber intensity after SERPINE1 overexpression (n = 30 cells).

### AP2A1 and SERPINE1 are reciprocally coupled in senescent fibroblasts

Given the role of AP2A1 in stress fiber regulation and integrin-associated trafficking [15], we next examined whether AP2A1 is functionally linked to SERPINE1 in senescent fibroblasts. In P31 fibroblasts, AP2A1 signal increased under stiffer substrate conditions and was accompanied by stronger stress fiber staining, suggesting that AP2A1 abundance is associated with stiffness-dependent cytoskeletal remodeling (Supplementary Fig. S5A,B).

Colocalization analysis using SIM imaging showed greater spatial overlap between AP2A1 and SERPINE1 in senescent cells than in young cells, suggesting a closer association between the two proteins in the senescent state (Figure 5A,B). To test whether AP2A1 influences SERPINE1 abundance, we perturbed AP2A1 expression in P31 cells. AP2A1 knockdown reduced intracellular SERPINE1 signal, whereas AP2A1 overexpression increased SERPINE1 signal (Figure 5C–F). We then examined the reciprocal relationship by manipulating SERPINE1 expression. SERPINE1 knockdown reduced AP2A1 signal, whereas SERPINE1 overexpression increased AP2A1 signal (Figure 5G–J). These reciprocal effects suggest that AP2A1 and SERPINE1 are functionally coupled in senescent fibroblasts.

**Figure 5.**
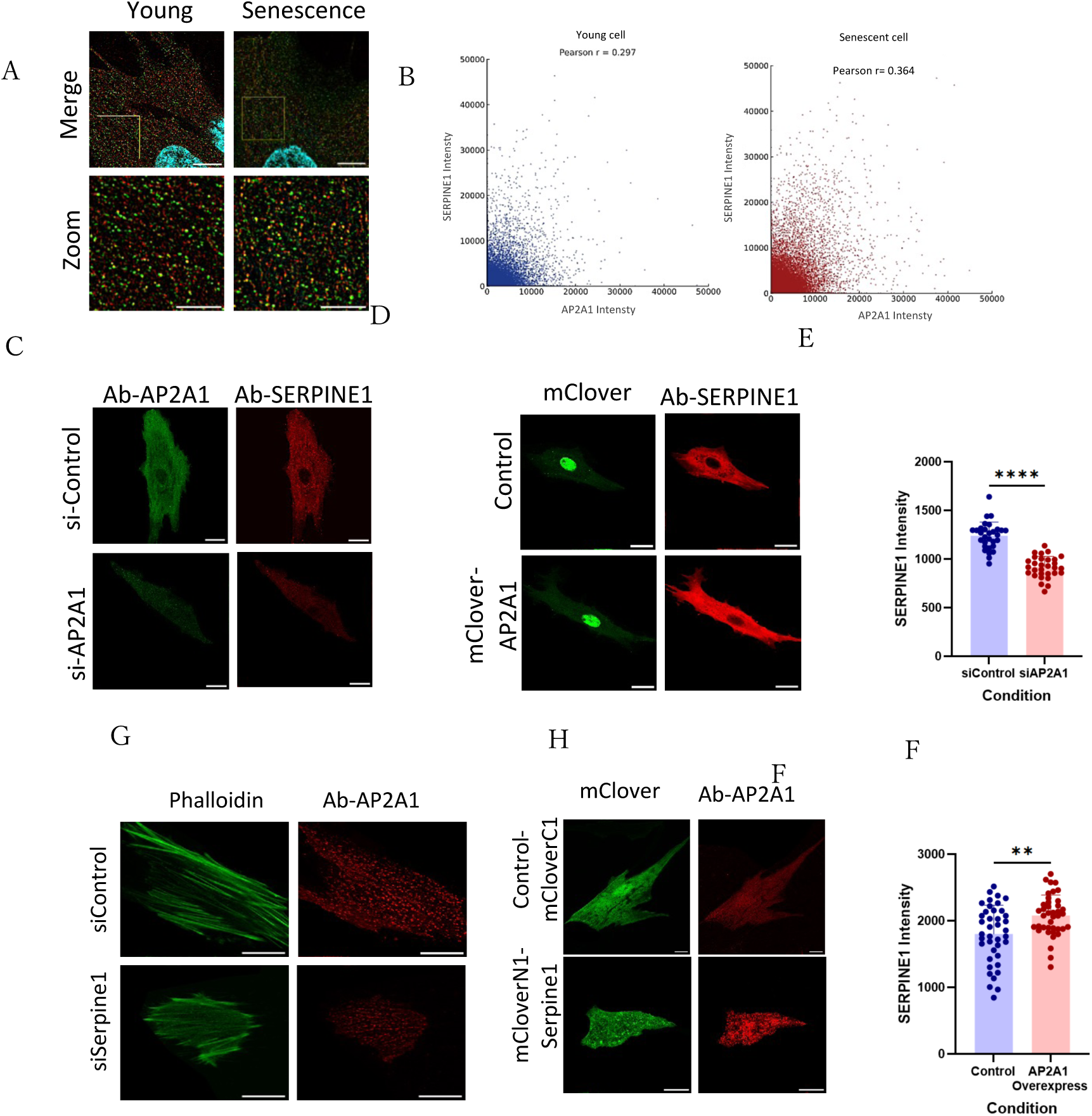
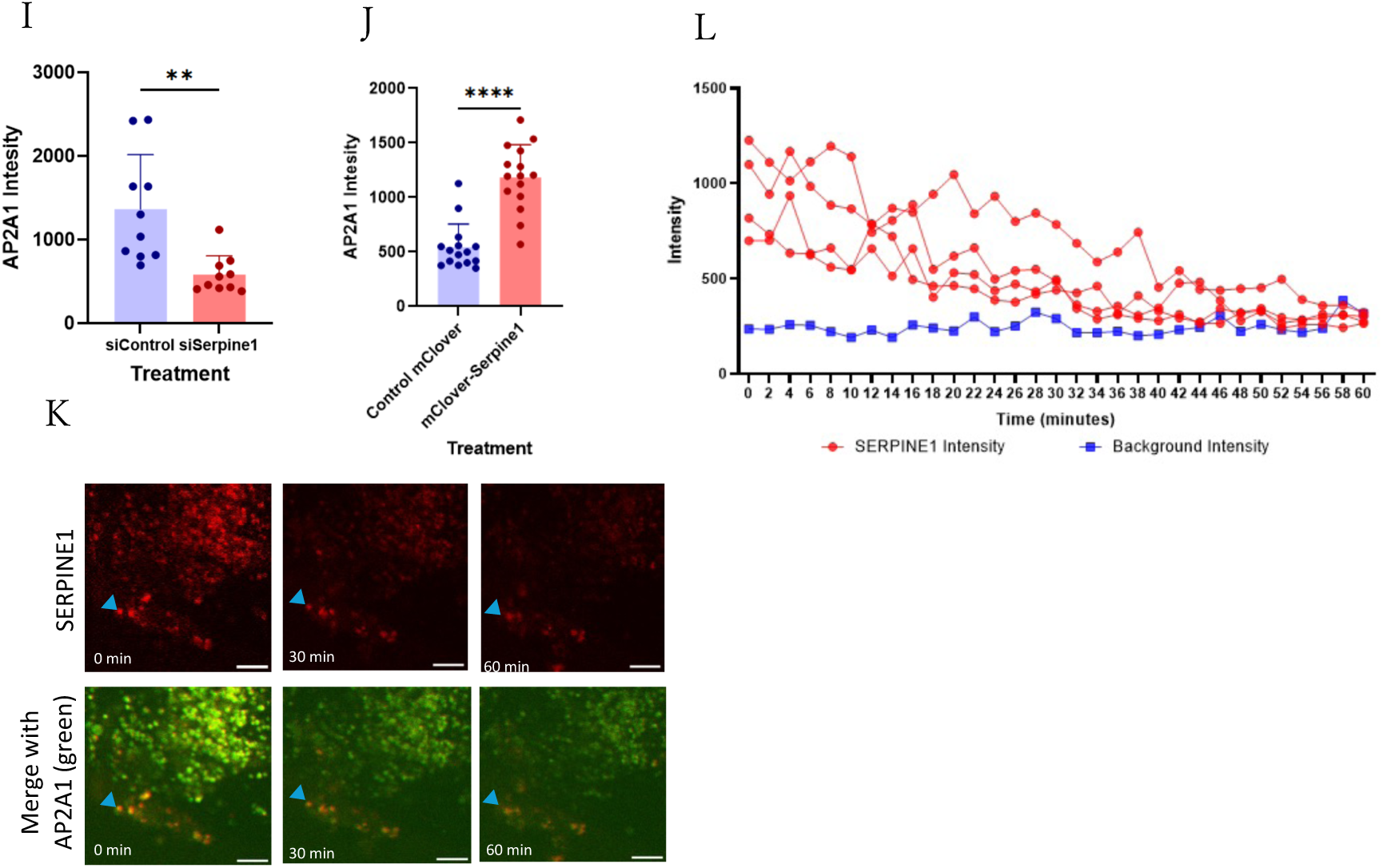
AP2A1 is functionally associated with SERPINE1 in senescent fibroblasts. A) Immunostaining showing colocalization of SERPINE1 (red) and AP2A1 (green) in young and senescent fibroblasts by structured illumination microscopy (SIM) (scale bar: 10 μm; inset: 5 μm). B) Pearson’s correlation coefficients for SERPINE1-AP2A1 colocalization in young and senescent fibroblasts (n = 3 cells per group). C) Immunostaining of SERPINE1 following AP2A1 knockdown (scale bar: 25 μm). D) Immunostaining of SERPINE1 following AP2A1 overexpression (scale bar: 25 μm). E) Quantification of SERPINE1 and AP2A1 fluorescence intensities after AP2A1 knockdown (n = 30 cells). F) Quantification of SERPINE1 fluorescence intensity after AP2A1 overexpression (n = 30 cells). G) Immunostaining of AP2A1 following SERPINE1 knockdown (scale bar: 25 μm). H) Immunostaining of AP2A1 following SERPINE1 overexpression (scale bar: 25 μm). I) Quantification of AP2A1 fluorescence intensity after SERPINE1 knockdown (n = 10 cells). J) Quantification of AP2A1 fluorescence intensity after SERPINE1 overexpression (n = 10 cells). K) Live-cell imaging of mCherry-SERPINE1 (red) and mClover-AP2A1 (green). The yellow triangle indicates a putative site of SERPINE1 secretion during time-lapse imaging (scale bar: 5 μm). L) Time-course analysis of fluorescence intensity at representative SERPINE1-positive regions and a background region.

Live-cell imaging revealed coordinated movement of mCherry-SERPINE1 with mClover-AP2A1 within senescent fibroblasts (Figure 5K). Time-course analysis showed that SERPINE1 signals tracked with AP2A1 over time, followed by selective disappearance of a subset of SERPINE1 signals while AP2A1 persisted in the same regions (Figure 5L). This sequence suggests a dynamic association between SERPINE1 and AP2A1 and is consistent with the disappearance of SERPINE1 from AP2A1-positive regions. However, because local disappearance of SERPINE1 signal does not distinguish intracellular dissociation from extracellular secretion, these observations do not directly define the route or mechanism of SERPINE1 secretion.

### Secreted SERPINE1 levels are altered by cellular senescence and SERPINE1/AP2A1 perturbation

Secreted SERPINE1/PAI-1 levels were measured in conditioned media by ELISA to assess extracellular SERPINE1 release during fibroblast senescence. After normalization to cell number, senescent fibroblasts showed significantly higher secreted SERPINE1 levels than young fibroblasts (Figure 7A). Additional analysis under different substrate conditions showed that senescence-associated differences in secreted SERPINE1 were context-dependent, with the clearest increase observed under rigid polystyrene culture conditions (Supplementary Fig. S6).

The effect of SERPINE1 and AP2A1 perturbation on SERPINE1 secretion was then examined in P31 senescent fibroblasts. SERPINE1 knockdown significantly reduced secreted SERPINE1 levels compared with control, confirming that the ELISA signal reflected SERPINE1 present in the conditioned medium (Figure 7B). AP2A1 knockdown showed a trend toward reduced secreted SERPINE1 levels; however, this difference did not reach statistical significance compared with the control (Figure 7B). Consistently, SERPINE1 overexpression strongly increased secreted SERPINE1 levels compared with control (Figure 7C). In contrast, AP2A1 overexpression did not significantly increase secreted SERPINE1 compared with control (Figure 7C). Together, these results indicate that extracellular SERPINE1 levels are strongly influenced by direct manipulation of SERPINE1 expression, whereas AP2A1 perturbation had comparatively modest effects.

## Discussion

Substrate stiffness promoted cell spreading in both young and senescent fibroblasts, but enhanced proliferation only in young cells. This divergence suggests that senescent fibroblasts retain mechanosensitive morphological responses while cell cycle progression remains constrained. In non-senescent contexts, ECM stiffening can promote fibroblast proliferation and activation through mechanosensitive pathways [24]. In contrast, senescent fibroblasts are limited by cell cycle arrest programs such as p16^INK4a^ and p21 [25], [26], [27], indicating that mechanical responsiveness and proliferative capacity become functionally decoupled. Thus, in senescent fibroblasts, extracellular mechanical cues are reflected primarily in cell spreading and cytoskeletal remodeling rather than proliferative expansion.

Our observation that substrate stiffness enhanced stress fiber thickening, focal adhesion enlargement, cellular stiffness, and senescence marker expression is consistent with the view that rigid extracellular environments reinforce mechanotransductive and senescence-associated programs in fibroblasts [21], [28]. Stress fibers and focal adhesions jointly regulate intracellular tension and cell–substrate force transmission; therefore, their coordinated enhancement on stiff substrates may provide a structural basis for the elevated mechanical phenotype observed in senescent fibroblasts [25], [29], [30]. Thus, matrix stiffening does not merely alter cell morphology but establishes a mechanically reinforced state in senescent fibroblasts.

SERPINE1/PAI-1 has been extensively studied as a senescence-associated secreted factor and matrix-remodeling mediator [17], [31], [32], and its expression is known to increase in response to stiff extracellular environments [33]. However, the intracellular organization and pre-secretory trafficking of SERPINE1 in senescent cells remain poorly understood. Our results address this gap by showing that substrate stiffness promotes stress fiber development, increases SERPINE1 abundance, and enhances the association of SERPINE1 with thickened stress fibers in senescent fibroblasts. Given the established roles of SERPINE1 in the SASP and fibrogenesis [34], [35], [36], its dual sensitivity to substrate stiffness and cellular senescence supports its role as a mediator linking mechanical remodeling to senescence-associated matrix dysfunction. Thus, SERPINE1 regulation in senescent fibroblasts is not limited to stiffness-dependent induction, but also involves its association with hypertrophic stress fibers characteristic of the senescent cytoskeletal state.

We propose that stiffness-dependent SERPINE1 regulation involves two interconnected routes (Figure 6). First, stiff substrates promote integrin-mediated adhesion, RhoA/ROCK-dependent stress fiber thickening, and focal adhesion maturation, thereby increasing intracellular tension [37], [38]. This tension-dependent state is known to activate YAP/TAZ signaling, which has been shown to cooperate with SMAD signaling to promote SERPINE1 transcription [33], [36], [39]. Although this pathway was not directly tested here, it provides a plausible route for the stiffness-dependent increase in SERPINE1 abundance observed in our experiments. Second, AP2A1-containing adaptor complexes, known regulators of clathrin-mediated trafficking, cargo sorting, and integrin-associated membrane dynamics [15], [40], are linked in our data to the organization of SERPINE1-containing compartments within stress fiber-rich regions. These observations support a role for AP2A1-positive, stress fiber-associated regions in SERPINE1 trafficking before secretion, although the precise secretion route remains unresolved.

**Figure 6.**
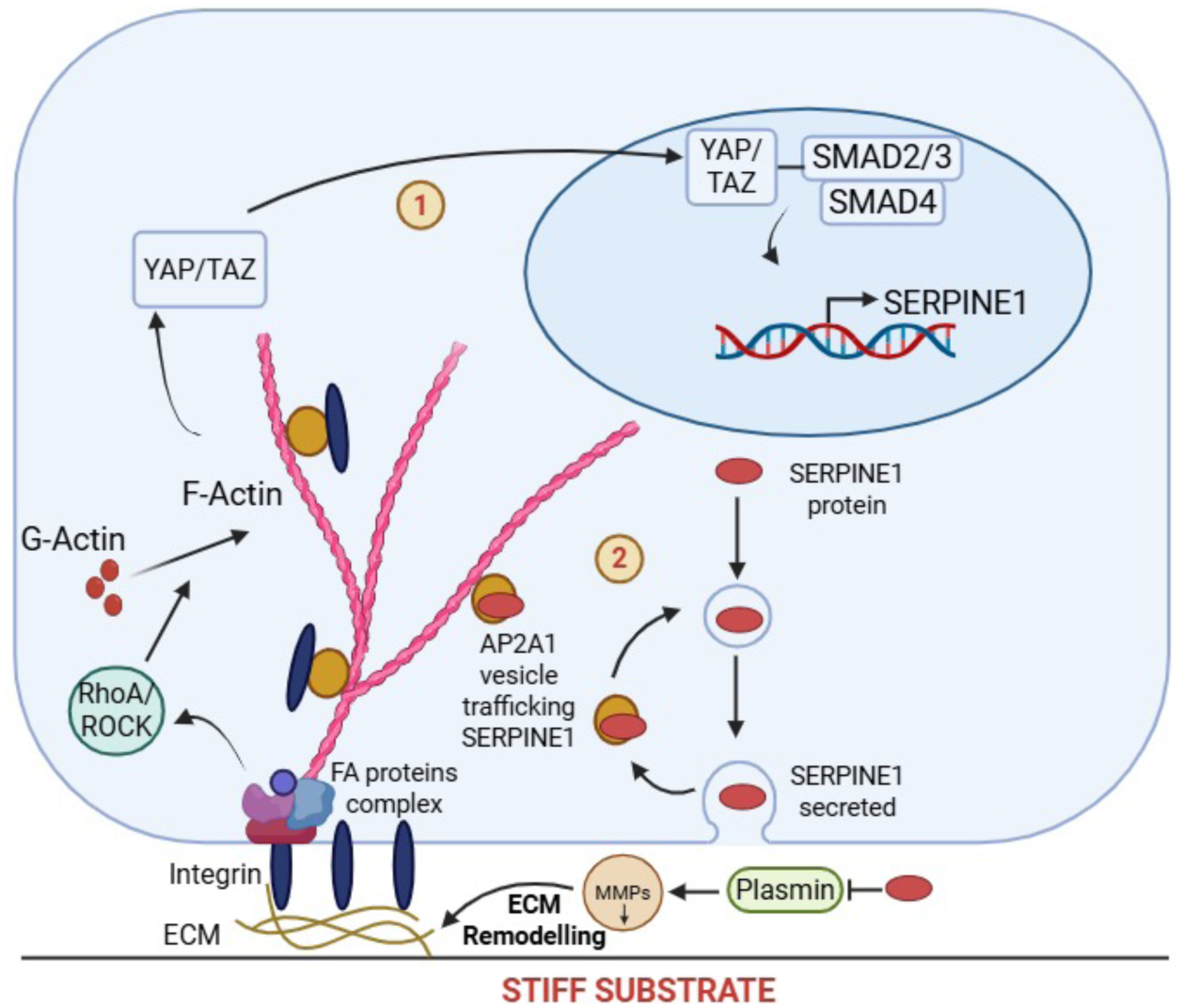
Proposed model linking extracellular stiffness, AP2A1, and SERPINE1 in senescent fibroblasts. Working model illustrating how extracellular stiffness and AP2A1-associated regulation may contribute to SERPINE1-associated cytoskeletal remodeling and ECM-related changes in senescent fibroblasts.

The reciprocal effects of SERPINE1 knockdown and overexpression indicate that SERPINE1 actively supports stress fiber organization in senescent fibroblasts, rather than serving only as a stiffness-responsive senescence marker. Consistently, previous studies have linked SERPINE1/PAI-1 to integrin signaling, RhoA/ROCK activity, actomyosin contractility [39], and ECM remodeling by limiting plasmin- and matrix metalloproteinase-dependent matrix degradation [41], [42], [43]. Reciprocal perturbation of AP2A1 and SERPINE1, together with their enhanced colocalization, coordinated movement, and selective loss of SERPINE1 signal from AP2A1-positive regions, further indicates that these proteins are functionally coupled rather than merely correlated. These observations support a mutually reinforcing regulatory module associated with the hypertrophic stress fiber network in senescent fibroblasts. This view is consistent with the established role of AP2A1-containing adaptor complexes in clathrin-mediated trafficking and membrane protein internalization [44], [45], as well as previous evidence linking AP2A1 to integrin trafficking and focal adhesion dynamics [15]. Because ventral stress fibers terminate at focal adhesions near the basal plasma membrane, this scaffold may help position SERPINE1-containing compartments in membrane-proximal regions, potentially favoring organized trafficking before extracellular release. Together, these findings place SERPINE1 as both a stiffness-responsive output and a functional contributor to the reinforced cytoskeletal state of senescent fibroblasts.

The ELISA data further support the extracellular relevance of SERPINE1 in senescent fibroblasts. Secreted SERPINE1/PAI-1 levels were higher in senescent fibroblasts after normalization to cell number, although additional substrate-dependent analysis indicated that this difference was most evident under rigid culture conditions, consistent with SASP-associated mechanosensitive secretion patterns [46], [47]. This suggests that extracellular SERPINE1 availability is associated with the senescent state, but may also depend on the mechanical context. Direct manipulation of SERPINE1 produced the clearest effect on secretion, with reduced secreted SERPINE1 after knockdown and increased secretion after overexpression, in line with studies demonstrating that SERPINE1 overexpression promotes senescence and ECM remodeling while its loss attenuates these effects [46], [48]. In contrast, AP2A1 perturbation produced weaker and statistically non-significant changes, suggesting that AP2A1 is unlikely to function as the sole determinant of SERPINE1 secretion. Instead, AP2A1 may act as a modulatory mediator that supports SERPINE1 availability through trafficking-associated organization, integrin-associated membrane dynamics, or cytoskeletal positioning of SERPINE1-containing compartments, consistent with the role of AP2-containing adaptor complexes in clathrin-mediated cargo sorting and integrin trafficking [15].

Once secreted, SERPINE1 inhibits plasmin generation and matrix metalloproteinase activation, thereby limiting ECM degradation and favoring matrix remodeling and stiffening [23], [49]. This established extracellular function provides a basis for a feedforward model in which substrate stiffness enhances SERPINE1 expression and trafficking, while SERPINE1-mediated matrix remodeling further supports mechanotransductive signaling. This model provides a framework for understanding how age-associated tissue stiffening may amplify fibroblast senescence and fibrosis-related remodeling, thereby promoting progressive tissue dysfunction. More broadly, our findings support the concept that ECM mechanical properties are not merely passive consequences of aging or fibrosis, but modifiable regulators of senescence-associated remodeling, highlighting their potential as therapeutic entry points for age-related tissue dysfunction.

**Figure 7.**
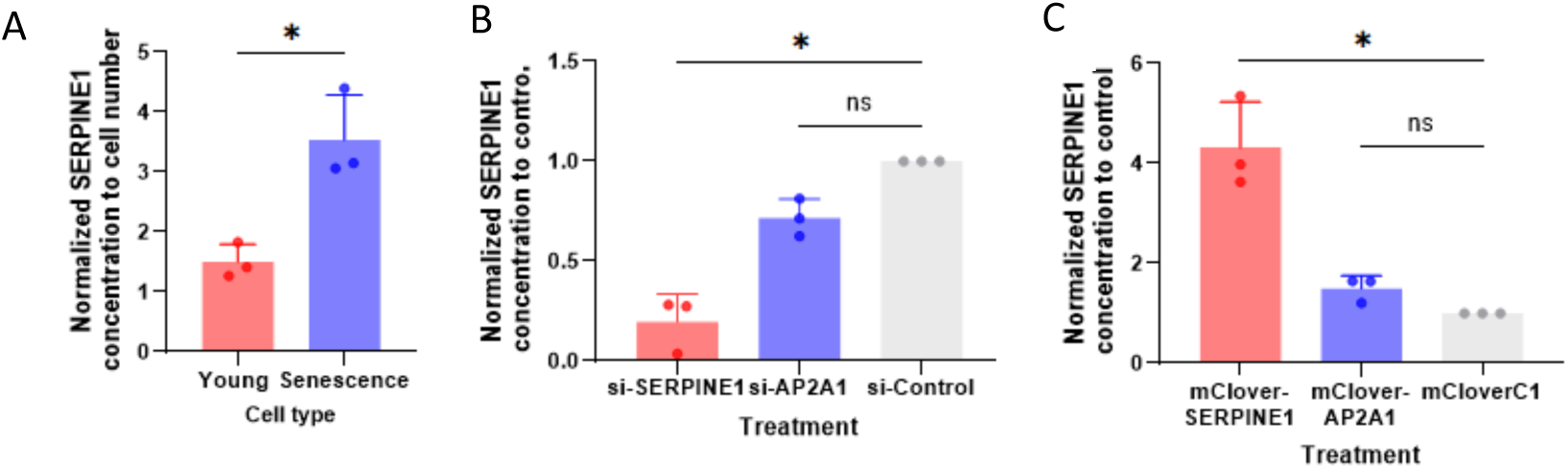
Secreted SERPINE1 levels after SERPINE1/AP2A1 perturbation and during cellular senescence. A) Secreted SERPINE1 concentration normalized to cell number in young and senescent fibroblasts. B) ELISA analysis of secreted SERPINE1 in conditioned medium after siRNA-mediated knockdown of SERPINE1 or AP2A1. C) ELISA analysis of secreted SERPINE1 after overexpression of mClover-SERPINE1 or mClover-AP2A1 (n = 3 independent experiments, * = p < 0.05; ns=not significant).

## Materials and methods

### Cell culture

Primary human foreskin fibroblasts (HFF-1, ATCC) were cultured in high-glucose Dulbecco’s Modified Eagle’s Medium (DMEM, Wako) supplemented with 15% fetal bovine serum (Sigma-Aldrich) and 1% penicillin-streptomycin (Wako). Cells were maintained at 37 °C with 5% CO₂ and sub-cultured at 1:3 when 70–80% confluent. Early-passage HFF-1 cells (P10–11) were used as the young group, whereas late-passage cells (P30–31) were used as the senescent group.

### Coverslip and gel preparation

Coverslips (ø 22 mm, Iwaki) were sterilized with 70% ethanol for 30 min, rinsed in sterile water, and air-dried. Coverslips were then activated with 2% APTES in absolute ethanol for 10 min, washed with ethanol and sterile water, and incubated with 0.5% glutaraldehyde in PBS for 30 min. After extensive rinsing, polyacrylamide hydrogels were prepared on activated coverslips using acrylamide:bis-acrylamide ratios of 8:0.48, 8:0.264, 5:0.3, and 5:0.1 to generate substrates of defined stiffness. Polymerization was initiated by ammonium persulfate (APS) and TEMED. After polymerization for 30–60 min, gels were coated with 0.2 mg/mL Sulfo-SANPAH in PBS, photoactivated under 365-nm UV light for 10 min, rinsed with PBS, and functionalized with fibronectin (10 µg/mL) for 1 h at 37 °C before cell seeding.

### Atomic force microscopy (AFM)

AFM measurements were performed using NanoWizard II (Osaka Univ.) BioAFM systems (JPK Instruments/Bruker) equipped with qp-BioAC-Cl-10 cantilevers (NANOSENSORS™) in PBS. Force–indentation curves were analyzed using the Hertz model to calculate Young’s modulus. AFM data were acquired and analyzed using JPK SPM Control Software.

### Cell growth rate and adhesion assay

Cell growth was quantified by manual counting using a pre-marked grid (0.2 × 0.2 mm) positioned beneath each culture surface. Ten fixed grid squares were selected per sample and imaged repeatedly at 24, 48, and 72 h. Cell counts at 24 h were used as the baseline for adhesion efficiency, and relative growth was calculated by normalizing cell numbers at 48 and 72 h to the 24-h value. Three independent gel preparations were analyzed for each stiffness condition.

### Edu proliferation assay

Cell proliferation was evaluated using the Click-iT EdU Imaging Kit (Alexa Fluor 647; Invitrogen) according to the manufacturer’s instructions. Cells were seeded on fibronectin-coated hydrogels and cultured for 24 h, followed by incubation with 20 µM EdU in culture medium for 24 h at 37 °C. Cells were then fixed with 3.7% paraformaldehyde for 15 min, permeabilized with 0.5% Triton X-100 in PBS for 20 min, and subjected to the Cu(I)-catalyzed Click-iT reaction for 30 min. Nuclei were counterstained with Hoechst 33342 (5 µg/mL in PBS). Samples were imaged using an Olympus FV3000 confocal laser scanning microscope equipped with a UPlan Apo 20× objective lens. The proliferation index was calculated as the percentage of EdU-positive nuclei relative to total Hoechst-positive nuclei.

### Cell area quantification

Cell area was measured using Fiji/ImageJ. The spatial scale was set using image metadata, and individual cell boundaries were manually outlined using the Polygon Selection tool. Cell area was quantified using the Measure function. For each condition, 30 cells were analyzed per experiment across three independent experiments.

### Quantification of cell stiffness and height

Cell stiffness was quantified by AFM as described above, with Young’s modulus calculated from force–indentation curves using the Hertz model. Cell height was determined from AFM topographic maps by measuring the maximum vertical distance between the substrate baseline and the apical cell surface. Five cells were analyzed per condition.

### Immunofluorescence staining and imaging

Cells cultured on glass coverslips or polyacrylamide hydrogels were fixed with 4% paraformaldehyde, permeabilized with 0.1% Triton X-100 in PBS, and blocked with 10% normal goat serum in PBS. Samples were incubated with primary antibodies against SERPINE1 (1:200; Santa Cruz Biotechnology) and AP2A1 (1:200; Thermo Fisher Scientific). Appropriate species-specific Alexa Fluor-conjugated secondary antibodies were used for detection, including goat anti-mouse Alexa Fluor 546 and goat anti-mouse Alexa Fluor 488. F-actin was stained with Alexa Fluor-conjugated phalloidin, and nuclei were counterstained with Hoechst 33342. Confocal images were acquired using an Olympus FV3000 microscope equipped with a UPlan Apo 60× oil-immersion objective (NA 1.42). Structured illumination microscopy (SIM) was performed using a Zeiss Elyra S1 system at the Center for Medical Innovation and Translational Research, Osaka University Medical School, equipped with an α Plan-Apochromat 100× oil-immersion objective (NA 1.46). Acquisition settings were kept constant within each experiment.

### Colocalization analysis

Colocalization between SERPINE1 and F-actin, and between SERPINE1 and AP2A1, was analyzed from confocal images (2048 × 2048 pixels) using the JACoP plugin in Fiji/ImageJ. Regions of interest (ROI) corresponding to individual cells were manually selected. Pearson’s correlation coefficient was used to quantify fluorescence colocalization between channels.

### SA-β-gal staining

Cellular senescence was assessed using a senescence-associated β-galactosidase staining kit (Cell Signaling Technology) following the manufacturer’s protocol. For this assay, young cells at P10 and senescent cells at P30 were cultured on substrates of varying gel stiffness in 12-well plates until reaching sub-confluence. After medium removal, cells were fixed with 4% paraformaldehyde in phosphate-buffered saline (PBS, Wako) at 37 °C, and subsequently incubated overnight with β-galactosidase staining solution (pH 6.0) in a dry incubator at 37 °C. Images were collected from 10 random fields using an inverted microscope (CKX41, Olympus) to determine the percentage of SA-β-gal–positive cells, identified by blue staining.

### Quantification of stress fiber thickness

The thickness of ventral stress fibers was quantified from grayscale images using Fiji/ImageJ. After background subtraction within each cell, a line region of interest (ROI) was drawn perpendicular to the width of an individual stress fiber, and the corresponding fluorescence intensity profile was generated. The positions of the two fiber edges were identified from the intensity profile, and stress fiber thickness was defined as the distance between these two edges. Approximately 300 measurements were analyzed in total.

### Western blotting

Whole-cell lysates were prepared from fibroblasts cultured under the indicated conditions by washing cells twice with cold PBS and scraping them into lysis buffer. Lysates were incubated on a shaker at 4 °C for 30 min and clarified by centrifugation at 15,000 × g for 20 min. Protein concentrations were determined using the Pierce BCA Protein Assay Kit (Thermo Fisher Scientific). Equal amounts of protein were separated by 10% SDS-PAGE and transferred onto 0.45-µm PVDF membranes (Wako). Membranes were blocked with 5% BSA in TBS-T for 1 h at room temperature and incubated overnight at 4 °C with primary antibodies against vinculin (1:5000; Wako), p53 (1:1000; Proteintech), and SERPINE1 (1:1000; Santa Cruz Biotechnology). After washing with TBS-T, membranes were incubated with HRP-conjugated secondary antibodies for 1 h at room temperature. Immunoreactive bands were visualized using the Immobilon Western Kit (Millipore), imaged with a ChemiDoc XRS+ system, and quantified using Image Lab software (Bio-Rad). For comparisons between young and senescent cells, band intensities were normalized to vinculin. For analyses across different substrate stiffness conditions, Ponceau S staining was used as the loading control.

### Plasmid construction and siRNA transfection

For SERPINE1 overexpression, the mClover2-SERPINE1 plasmid was generated by inserting a synthetic human SERPINE1 coding sequence into the mClover2-N1 backbone (Addgene) using restriction enzyme cloning with AgeI and NheI. The mCherry-SERPINE1 construct was generated by Gibson assembly using fragments derived from mClover2-SERPINE1 and cloned into an mCherry backbone. The mClover2-AP2A1 construct was prepared as previously described [15]. All plasmids were sequence-verified before use.

For knockdown experiments, siRNAs targeting SERPINE1 (WD10956708 and WD10956709; Sigma-Aldrich) and AP2A1 (s183 and s184; Thermo Fisher Scientific) were transfected at 30 pmol. A negative control siRNA was obtained from Cell Signaling Technology. Plasmid transfections were performed using Lipofectamine LTX with Plus Reagent (Invitrogen) with 2.5 µg DNA per 35-mm dish at 50–70% confluence in Opti-MEM. siRNA transfections were performed using Lipofectamine RNAiMAX (Invitrogen) in Opti-MEM. Cells were analyzed 24 h after plasmid transfection and 48 h after siRNA transfection.

### Time-lapse imaging of SERPINE1 and AP2A1

Cells at P30 were cotransfected with mClover2-AP2A1 (green) and mCherry-SERPINE1 (red). Simultaneous time-lapse imaging of AP2A1 and SERPINE1 was performed using confocal microscopy (FV3000) at 2-min intervals for 1 h. Protein movement was analyzed using the TrackMate plugin in Fiji/ImageJ. A total of 5 cells were analyzed, and 10 representative protein trajectories were tracked per cell.

### Secreted SERPINE1/PAI-1 quantification

Secreted SERPINE1/PAI-1 levels were measured in conditioned media using a Human PAI-1 ELISA Kit (Invitrogen, Thermo Fisher Scientific, BMS2033) according to the manufacturer’s instructions. Young, senescent, and perturbed P31 senescent fibroblasts were washed with PBS and incubated in serum-free DMEM for 24 h before media collection. Conditioned media were centrifuged to remove cell debris, and SERPINE1 concentrations were calculated from the standard curve and normalized to cell number. Perturbation data were expressed relative to control.

### Statistical analysis

All experiments were independently repeated at least three times unless otherwise stated. Data are presented as mean ± standard deviation (SD). Statistical analyses were performed using GraphPad Prism version 10. Comparisons among multiple groups were analyzed using one-way ANOVA followed by Tukey’s multiple-comparison test. Comparisons between two groups were performed using an unpaired two-tailed Student’s t-test. Statistical significance was defined as p < 0.05 (*), p < 0.01 (**), p < 0.001 (***), and p < 0.0001 (****).

## Acknowledgments

This study was partly supported by JSPS KAKENHI grant (23H04928 and 25K03456).

## Author contributions

Conceptualization: I.C.N., S.D.; Formal analysis: I.C.N.; Investigation: I.C.N.; Resources: P.C., T.S., C.B., S.D.; Writing - original draft: I.C.N.; Writing - review & editing: S.D.; Funding acquisition: S.D.

## Data availability

All relevant data can be found within the article and its supplementary information.

---

**Supplementary Figure S1.**
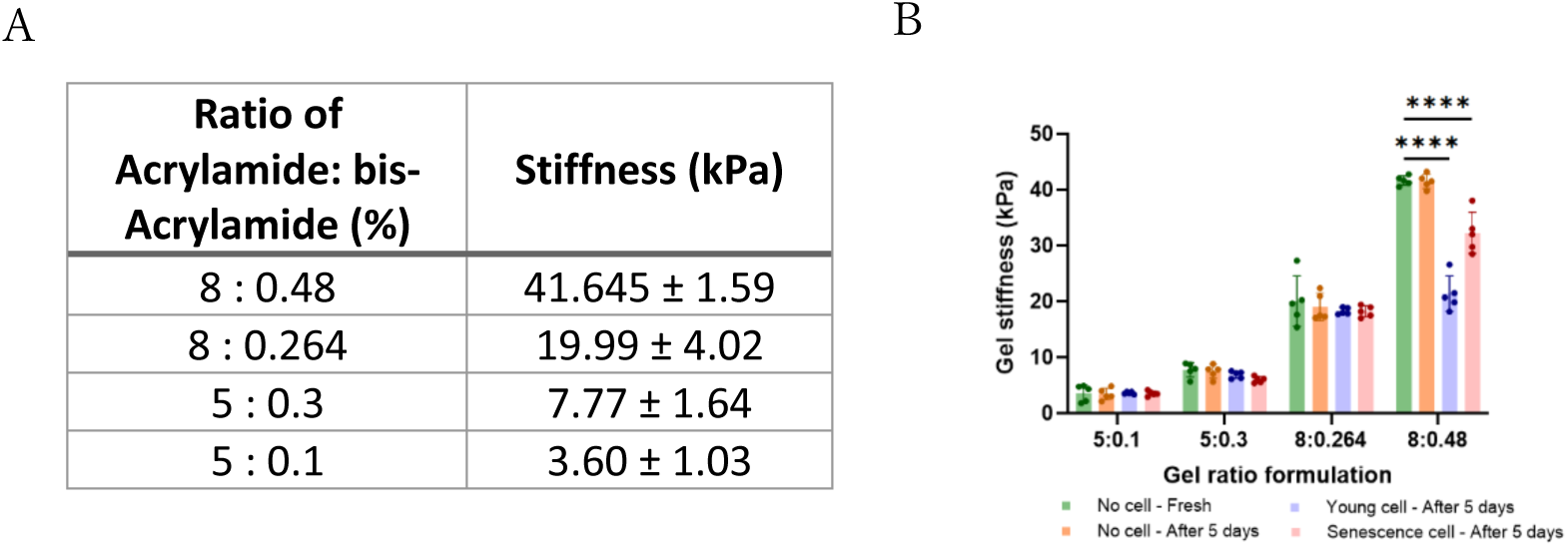
Polyacrylamide hydrogel validation and stability. A) Young’s modulus measurement (AFM) of hydrogels of different acrylamide/bis-acrylamide ratios. B) Assessment of gel stiffness stability over 5 days at 37°C (n = 5 per group)

**Supplementary Figure S2.**
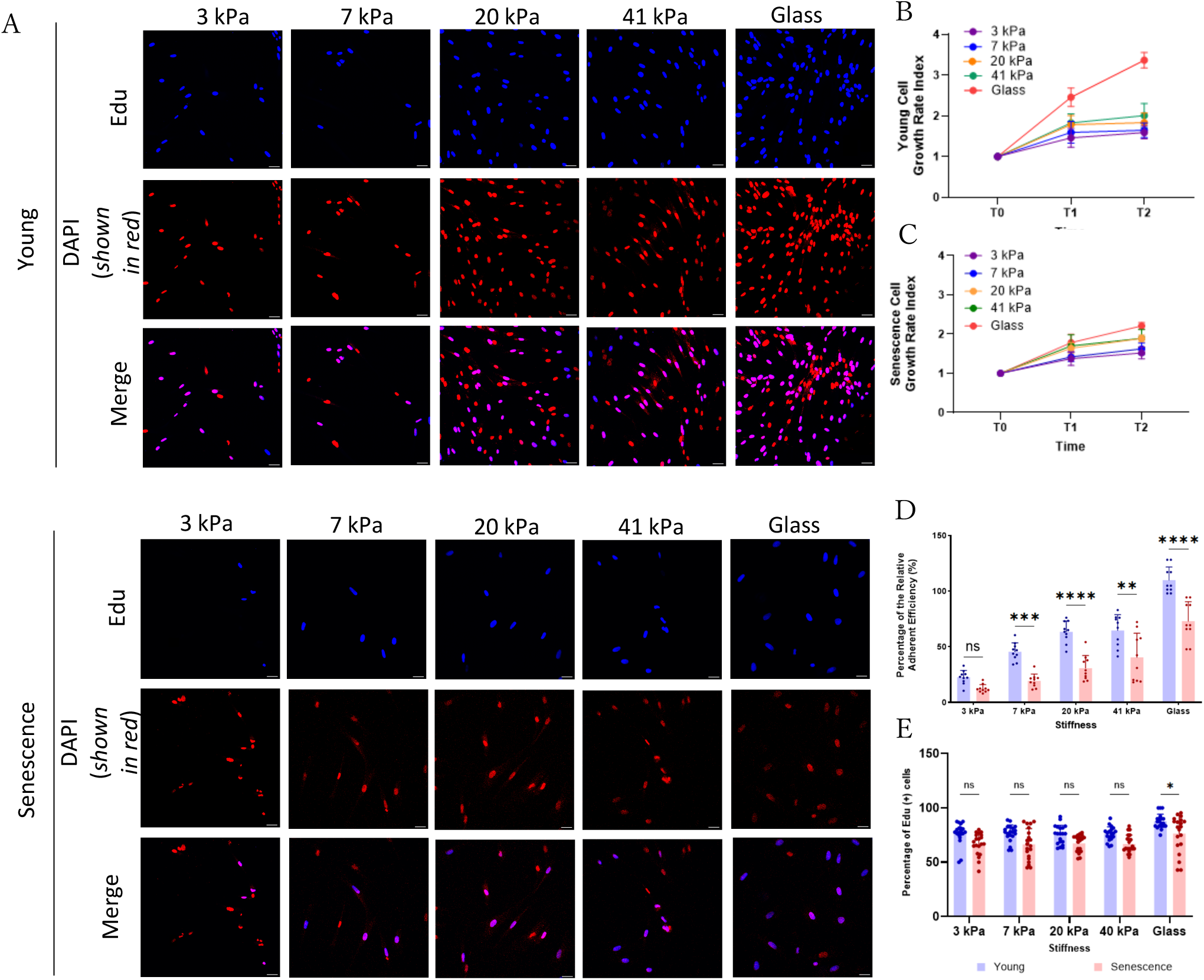
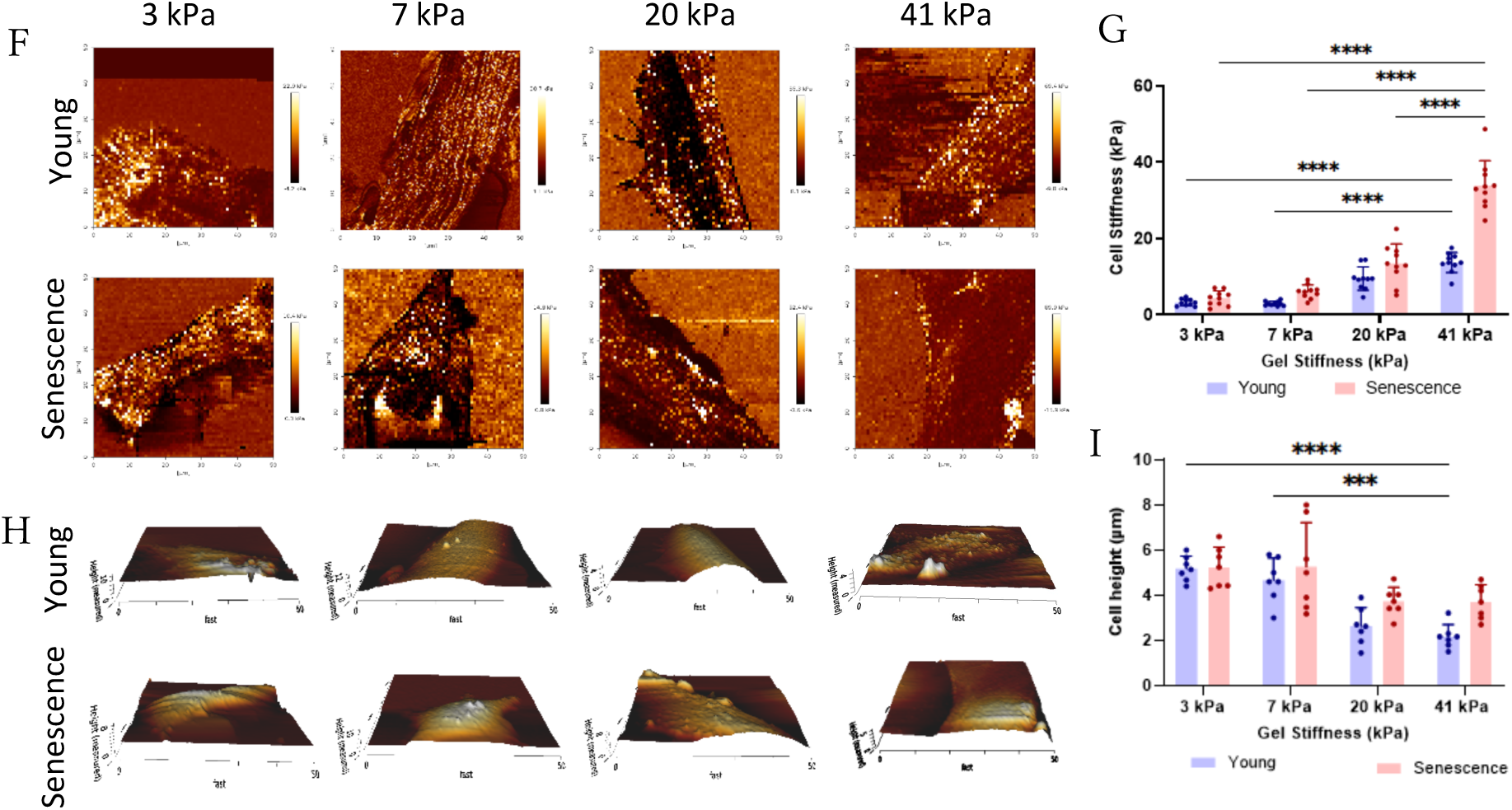
Mechanical profiling of fibroblasts on hydrogels. A) Representative immunofluorescence images of young fibroblasts cultured on hydrogels of 3, 7, 20, 41 kPa and glass, stained with EdU (green) and DAPI (red). Merged images show proliferating nuclei (20×, scale bar: 50 µm, n = 20). B-C) Quantification of the growth rate of young cells (B) and senescent cells (C) on various substrate stiffness (n = 10). D) The quantification percentage of relative adhesion after 24h incubation. E) EdU proliferation assay images and quantification for young and senescent fibroblasts on different stiffnesses (n = 20; scale bar, 50 μm). F-G) AFM topography (F) and quantification (G) of cellular stiffness (Young’s modulus) across hydrogels of different stiffness (n = 5 cells). H-I) AFM topography (H) and quantification (I) of maximum cell height across stiffness conditions (n = 5 cells).

**Supplementary Figure S3.**
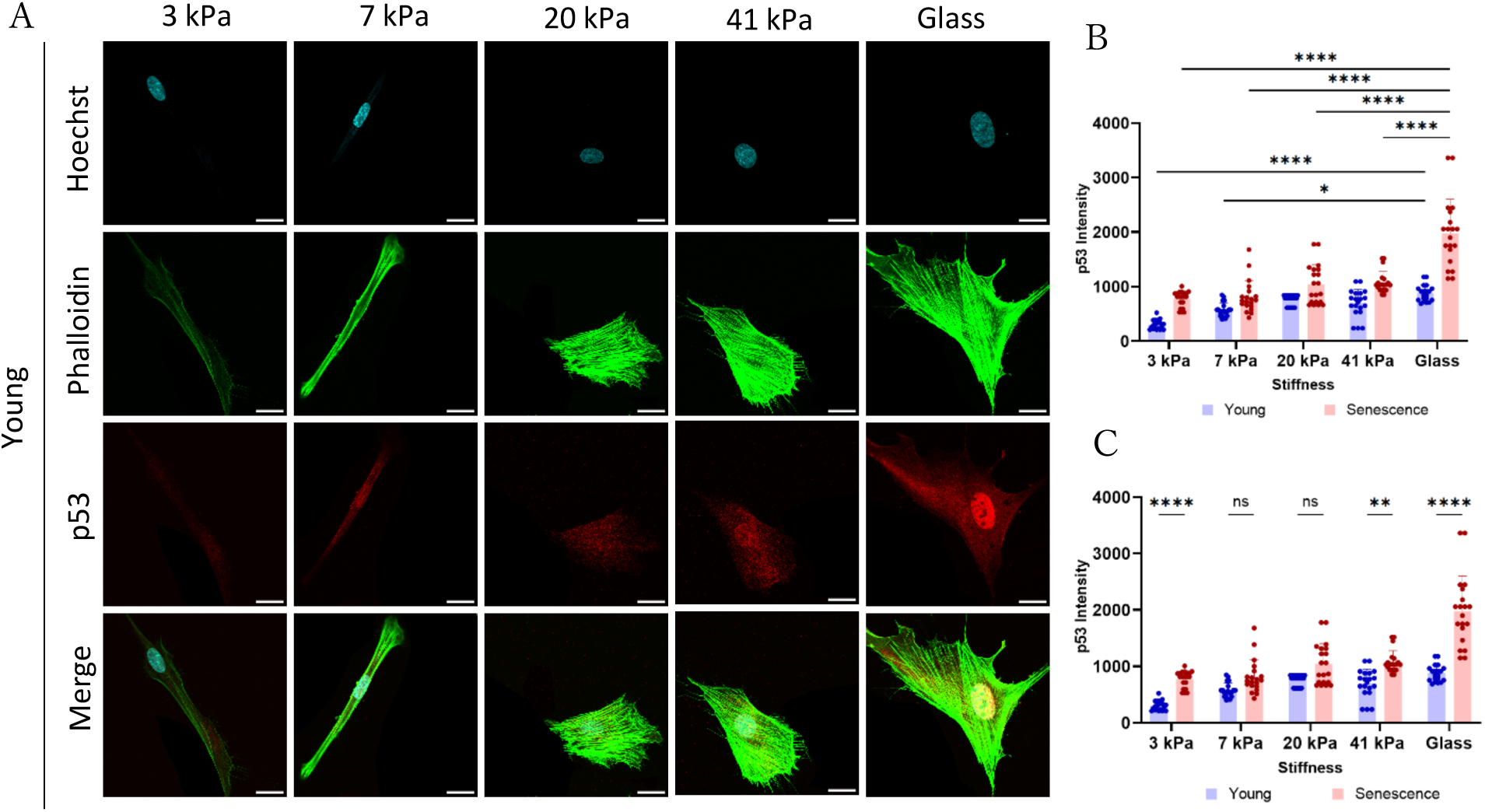
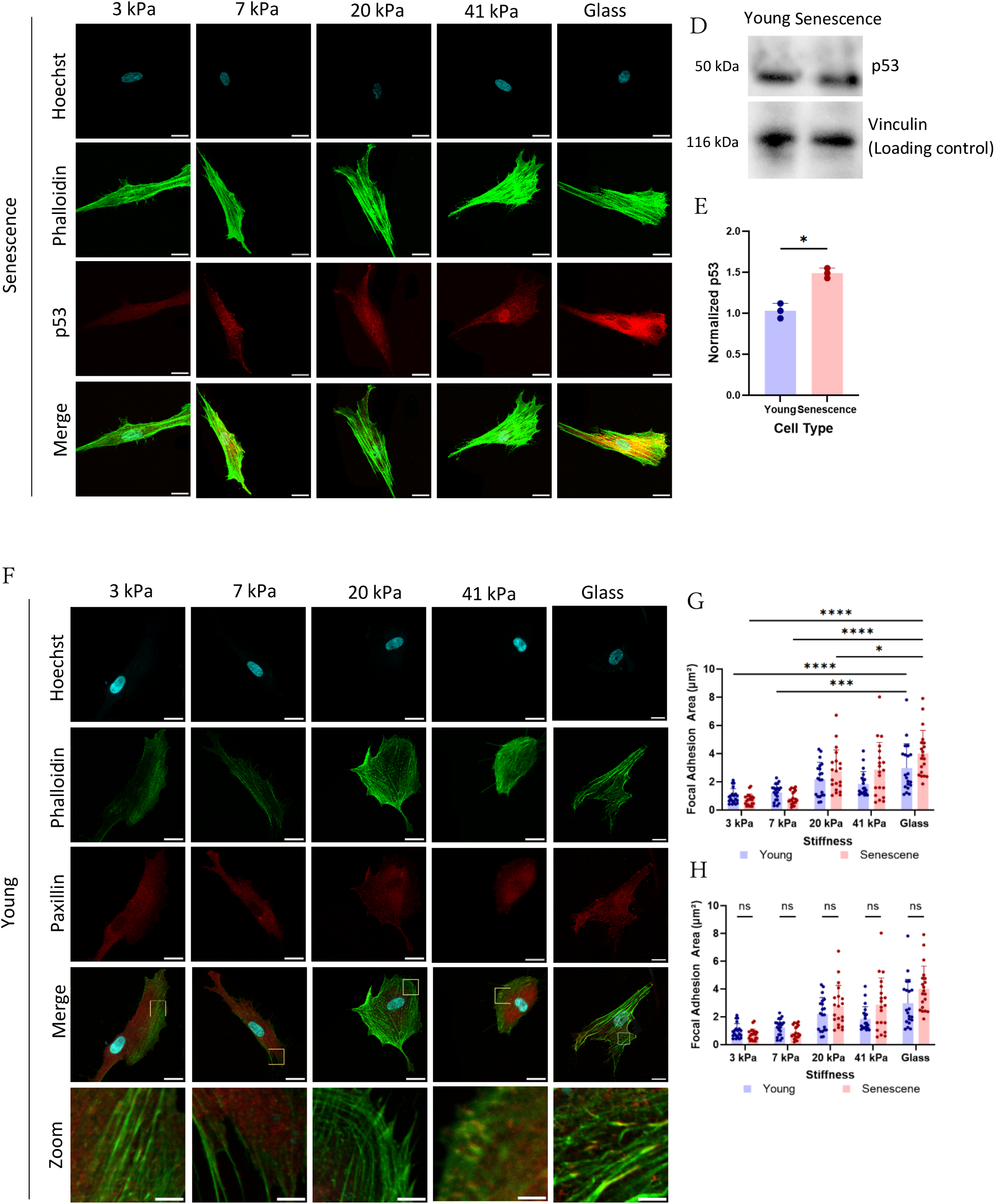
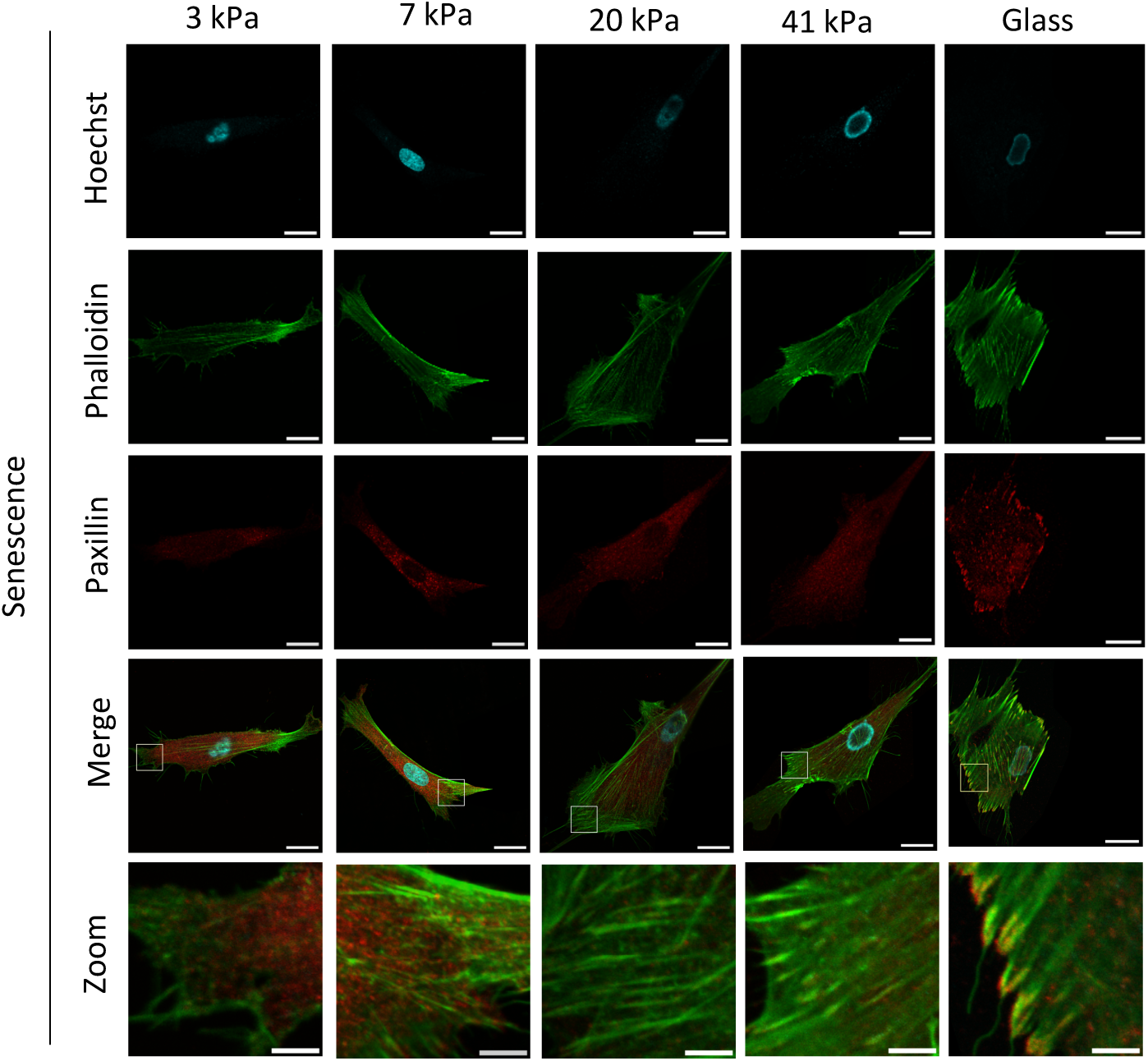
p53 expression and focal adhesion responses across substrate stiffness in young and senescent fibroblasts. A) Immunofluorescence staining of fibroblasts cultured on 3, 7, 20, 41 kPa hydrogels and glass (Scale bar: 25 µm). B-C) p53 intensity comparison in various hydrogel stiffnesses. Statistical comparison was performed between treatments and control (B) and between young and senescent cells (C) (n = 30 cells). D) Western blot of p53 expression in young and senescent fibroblasts, vinculin as loading control. E) Quantification of p53 protein normalized to vinculin as control (n = 3 cells). F) Immunofluorescence staining of paxillin-labeled focal adhesions in young and senescent fibroblasts. Zoomed insets highlight focal adhesion morphology (Scale bar: 25 μm, zoom: 5 μm). G-H) Quantification of focal adhesion area across substrate stiffness conditions. Statistical comparisons were performed between each stiffness condition and the control group (G) and between young and senescent cells at each stiffness (H) (n = 50).

**Supplementary Figure S4.**
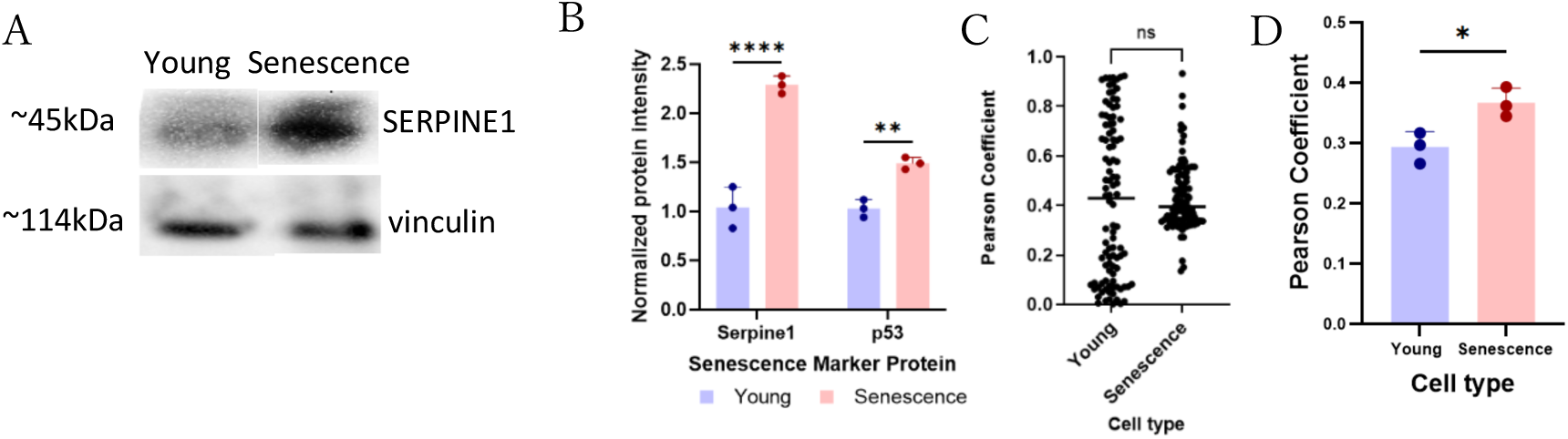
SERPINE1 expression and colocalization analysis in young and senescent fibroblasts. (A) Western blot comparison of SERPINE1 expression in lysates from young and senescent fibroblasts. Vinculin was used as the loading control. (B) Quantification of SERPINE1 band intensity normalized to vinculin from three biological replicates. (C) Quantification of SERPINE1 colocalization with actin stress fibers (Pearson’s correlation coefficient). (D) Quantification of SERPINE1 colocalization with AP2A1 (Pearson’s correlation coefficient).

**Supplementary Figure S5.**
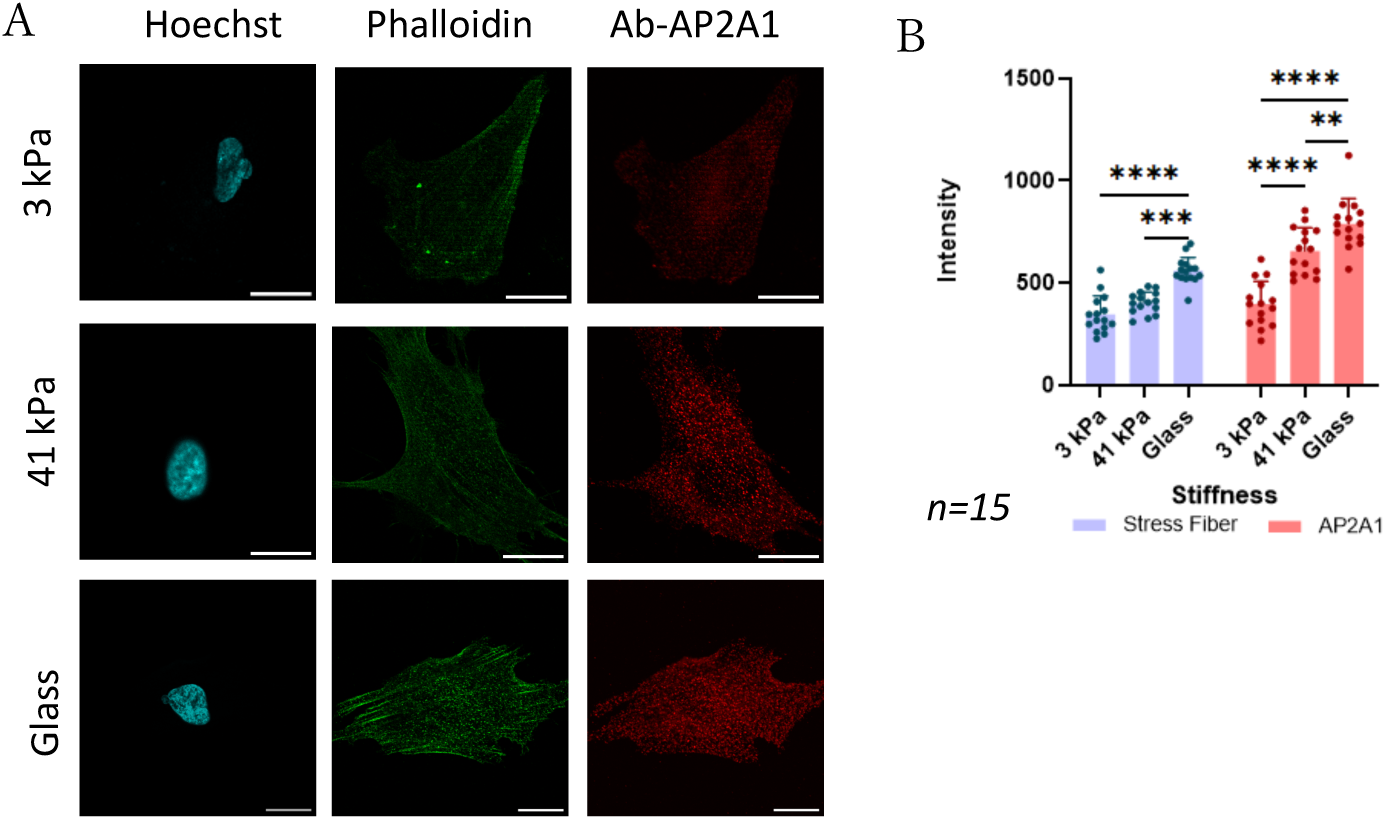
AP2A1 expression across soft (3 kPa), stiff (41 kPa), and glass substrates in senescent human fibroblasts. (A) Representative immunofluorescence images of AP2A1 (red), F-actin/stress fibers (green), and nuclei (blue) in senescent fibroblasts cultured on 3 kPa, 41 kPa, and glass substrates. (B) Quantification of AP2A1 and stress fiber intensities (n = 15 cells).

**Supplementary Figure S6.**
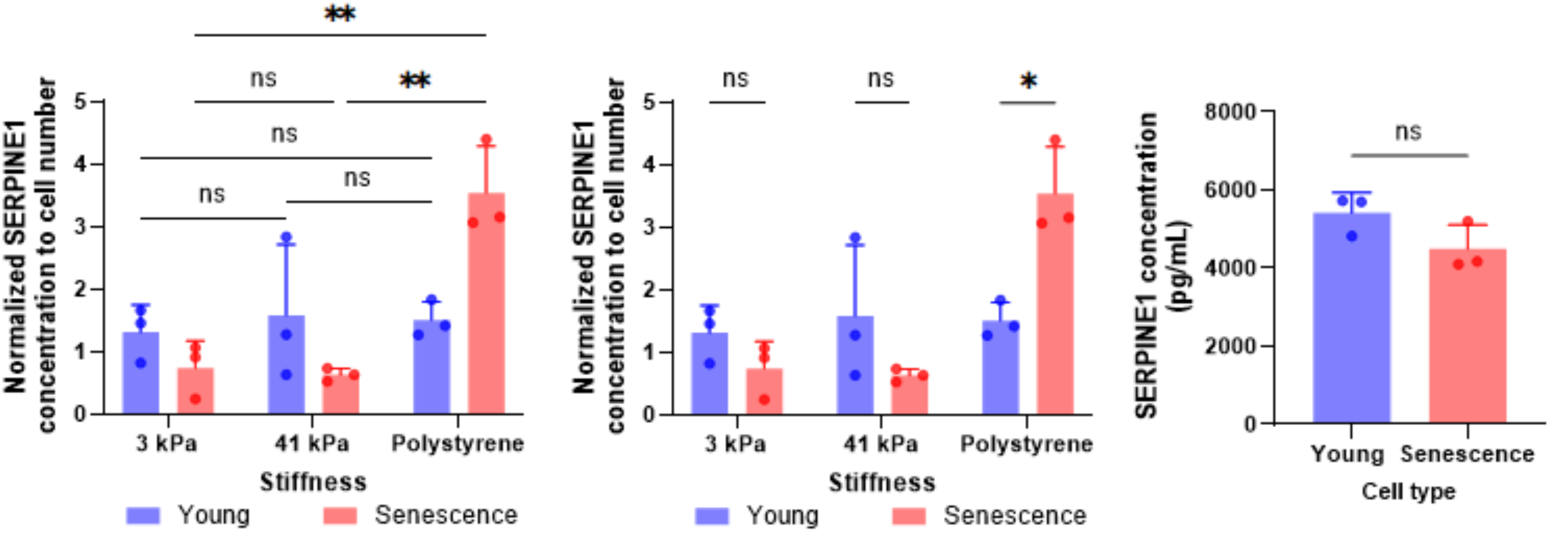
Substrate stiffness-dependent SERPINE1 secretion in young and senescent fibroblasts. A) Secreted SERPINE1/PAI-1 levels in conditioned medium from young and senescent fibroblasts cultured on 3 kPa, 41 kPa, and polystyrene substrates. SERPINE1 concentration was normalized to cell number. B) Comparison of normalized secreted SERPINE1/PAI-1 levels between young and senescent fibroblasts under each substrate condition. C) Basal SERPINE1/PAI-1 concentration in conditioned medium from young and senescent fibroblasts. Data are presented as mean ± SD; n = 3 independent experiments; *p < 0.05; **p < 0.01; ns, not significant.

